## Supplementary Table 2 for "A meta-analysis of the inhibin network reveals prognostic value in multiple solid tumors"

| **Type** | | **Subtype** | **Variable** | ***INHA*** | ***INHBA*** | ***INHBB*** | ***TGFBR3*** | ***Eng*** |
| --- | --- | --- | --- | --- | --- | --- | --- | --- |
| **Breast Cancer^#^** | | **All** | p value | **5.9E-8** | .29 | **.034** | **2.2E-12** | **.0014** |
|  |  |  | Hazard Ratio (HR) | **.74** | 1.06 | **.89** | **.69** | **.84** |
|  |  | **p53 Mutated** | p value | **.0056** | .2 | .3 | .41 | .12 |
|  |  |  | Hazard Ratio (HR) | **1.99** | 1.37 | 1.29 | .82 | 1.46 |
|  |  | **TNBC** | p value | 0.068 | .064 | .9 | .061 | .68 |
|  |  |  | Hazard Ratio (HR) | 1.49 | 1.5 | 1.03 | .67 | 1.09 |
|  |  | **Luminal A** | p value | **.00021** | .95 | **.0014** | **1.5E-6** | .23 |
|  |  |  | Hazard Ratio (HR) | **.72** | 1.01 | **.76** | **.66** | .9 |
|  |  | **Luminal B** | p value | **5.9E-5** | .33 | .78 | .076 | **.0057** |
|  |  |  | Hazard Ratio (HR) | **.67** | 1.1 | .97 | .84 | **.76** |
|  |  | **HER2+** | p value | .25 | .65 | .54 | .14 | .2 |
|  |  |  | Hazard Ratio (HR) | .8 | .92 | 1.13 | .75 | .78 |
| **Serous Ovarian Cancer*** | | **All** | p value | **1.5E-6** | **.047** | .16 | .096 | **.0032** |
|  |  |  | Hazard Ratio (HR) | **.71** | **1.16** | .9 | 1.18 | **.82** |
|  |  | **p53 Mutated** | p value | **.00039** | **.0055** | .079 | **.039** | **.0098** |
|  |  |  | Hazard Ratio (HR) | **1.55** | **1.42** | 1.23 | **.79** | **1.36** |
| **Lung Cancer** | | **All** | p value | **.00029** | .37 | .78 | **3.4E-7** | **.0056** |
|  |  |  | Hazard Ratio (HR) | **1.26** | .94 | 1.02 | **.65** | **1.2** |
|  |  | **Adenocarninoma** | p value | **5.6E-9** | .59 | .18 | **5.5E-10** | **1.6E-8** |
|  |  |  | Hazard Ratio (HR) | **2.01** | 1.07 | 1.17 | **.46** | **1.98** |
|  |  | **Squamous Cell Carcinoma** | p value | .14 | .26 | .67 | .97 | .79 |
|  |  |  | Hazard Ratio (HR) | 1.2 | .87 | 1.05 | 1.01 | 1.03 |
| **Gastric Cancer** | | **All** | p value | **3.2E-8** | **.0053** | **2E-15** | **.019** | **2E-9** |
|  |  |  | Hazard Ratio (HR) | **1.66** | **1.31** | **2.17** | **1.23** | **1.78** |
|  |  | **HER2-** | p value | **.00055** | .21 | **5.3E-13** | **.0013** | **8E-7** |
|  |  |  | Hazard Ratio (HR) | **1.53** | 1.17 | **2.64** | **1.48** | **1.75** |
|  |  | **HER2+** | p value | **.0029** | **2.2E-5** | **7.6E-5** | **.035** | **.0032** |
|  |  |  | Hazard Ratio (HR) | **1.51** | **1.78** | **1.69** | **1.33** | **1.62** |
| **Cervical Cancer** | **Squamous Cell Carcinoma** | | p value | **.022** | **1.5E-5** | .19 | .35 | .12 |
|  |  |  | Hazard Ratio (HR) | **1.97** | **2.94** | 1.38 | 1.32 | 1.49 |
| **Head and Neck Cancer** | **Squamous Cell Carcinoma** | | p value | **.035** | **0.00044** | **0.0037** | **.00049** | **.0022** |
|  |  |  | Hazard Ratio (HR) | **1.42** | **1.68** | **1.52** | **.62** | **.64** |
| **Kidney Cancer** | **Renal Clear Cell Carcinoma** | | p value | **7.1E-06** | .19 | .32 | **2.1E-7** | **8.6E-6** |
|  |  |  | Hazard Ratio (HR) | **1.98** | .82 | 1.16 | **.46** | **.51** |
|  | **Renal Papillary Cell Carcinoma** | | p value | **.0062** | **.00011** | **.0056** | **.042** | .091 |
|  |  |  | Hazard Ratio (HR) | **2.24** | **3.16** | **2.33** | **.53** | 1.68 |
| **Liver Cancer** | **Hepatocellular Carcinoma** | | p value | .38 | **.02** | .093 | **.021** | **5.6E-6** |
|  |  |  | Hazard Ratio (HR) | 1.17 | **1.52** | 1.39 | **.67** | **.45** |
| **Endometrial**  **Cancer** | **Uterine Corpus** | | p value | **.046** | .11 | **.011** | **.01** | **.025** |
|  |  |  | Hazard Ratio (HR) | **.65** | .71 | **.49** | **1.74** | **.54** |
| **Brain Cancer** | **Glioblastoma** | | p value | **.019** | .94 | .2767 | .832 | .318 |
|  |  |  | Hazard Ratio (HR) | **1.27** | 1.08 | 1.02 | 1.085 | .806 |
|  | **Low Grade Glioma** | | p value | **.005** | .6925 | **.046** | .7413 | **.0004** |
|  |  |  | Hazard Ratio (HR) | **.7088** | .955 | **1.208** | .9907 | **1.675** |
